# Single-cell RNA sequencing of mutant whole mouse embryos: from the epiblast to the end of gastrulation

**DOI:** 10.1101/2024.04.29.591777

**Authors:** Elizabeth Abraham, Mikel Zubillaga, Thomas Roule, Eleonora Stronati, Naiara Akizu, Conchi Estaras

## Abstract

Over the last decade, single-cell approaches have become the gold standard for studying gene expression dynamics, cell heterogeneity, and cell states within samples. Before single-cell advances, the feasibility of capturing the dynamic cellular landscape and rapid cell transitions during early development was limited. In this paper, we designed a robust pipeline to perform single-cell and nuclei analysis on mouse embryos from E6.5 to E8, corresponding to the onset and completion of gastrulation. Gastrulation is a fundamental process during development that establishes the three germinal layers: mesoderm, ectoderm, and endoderm, which are essential for organogenesis. Extensive literature is available on single-cell omics applied to WT perigastrulating embryos. However, single-cell analysis of mutant embryos is still scarce and often limited to FACS-sorted populations. This is partially due to the technical constraints associated with the need for genotyping, timed pregnancies, the count of embryos with desired genotypes per pregnancy, and the number of cells per embryo at these stages. Here, we present a methodology designed to overcome these limitations. This method establishes breeding and timed pregnancy guidelines to achieve a higher chance of synchronized pregnancies with desired genotypes. Optimization steps in the embryo isolation process coupled with FAST genotyping protocol (3 hours) allow for microdroplet-based single-cell to be performed on the same day, ensuring the high viability of cells and robust results. We also include guidelines for optimal nuclei isolations from embryos. Thus, these approaches increase the feasibility of single-cell approaches of mutant embryos at the gastrulation stage. We anticipate this method will facilitate the analysis of how mutations shape the cellular landscape of the gastrula.

**SUMMARY:** We establish a pipeline for high-quality single-cell and nuclei suspensions of gastrulating mouse embryos for sequencing of single cells and nuclei.

## INTRODUCTION

Gastrulation is a fundamental process required for normal development. This rapid and dynamic process occurs when pluripotent cells transition into lineage-specific precursors that define how our organs form. For years, gastrulation was long defined as the formation of three largely homogeneous populations: mesoderm, ectoderm, and endoderm. However, high-resolution technologies and an emerging number of embryonic stem cell models^1,2^ unveil an unprecedented heterogeneity among the early germ layers^3,4^. This suggests that much more remains to be uncovered about the mechanisms regulating the distinct cell populations of the gastrula. Mouse embryonic development has been one of the best models to study early cell fate decisions during gastrulation^3,5^. Gastrulation in mice is rapid as the entire process of gastrulation occurs within 48 hours, from embryonic day E6.5 to E8^5^.

Recent advancements in single-cell technologies have enabled detailed mapping of wild-type mouse embryonic development, providing a comprehensive overview of the cellular and molecular landscapes of embryos during gastrulation^3,4,6,7,8^. However, the analysis of mutant embryos at these stages is less common and often limited to FACS-sorted populations^9,10^. The scarce literature reflects the technical challenges associated with the manipulation and single-cell preparation of gastrulating embryos that require genotyping. In this study, we provide a strategy designed to obtain robust single-cell data from mutant gastrulating embryos from E6.5 to E8. Our strategy is designed to overcome constraints associated with the low number of embryos with desired genotype per pregnancy, and the decrease in viability caused by freezing-thawing embryos or cells.

Here, we describe an optimized methodology from the establishment of timed pregnancies to final sequencing of single-cells/nuclei. Our method explains how to increase the number of synchronized pregnancies to obtain a higher number of embryos with desired genotype, cell/nuclei isolations to improve the viability of the cells, and a FAST-genotyping protocol. We also describe the process of embryo isolation at different gastrulation timepoints. Our methodology helps to increase the number of final viable embryo cells/nuclei for sequencing, to ensure high-quality sequencing data. Therefore, we believe that this method will open the doors for single-cell studies of gastrulating embryos that require genotyping. Our method can be applied to genotyping for biological sex, often required in works supported by NIH-funding.

## PROTOCOL

This protocol and all animal experiments described were formally approved and in accordance with institutional guidelines established by the Temple University Institutional Animal Care and Use Committee, which follows the Association for Assessment and Accreditation of Laboratory Animal Care international guidelines. All mice described were on the C57/BL6N background strain. No animal health concerns were observed in these studies.

### 1 Breeding Colony and Timed Pregnancies

1.1 Pregnancies will be timed by the visualization of a vaginal plug. The noon of the day of the plug is considered E0.5.
1.2 House mice using general husbandry practices. Track and collect colony information every day when timed breeding is occurring. Log all information such as the age of the mouse, the number of pregnancies a female mouse has had, the number of plugs placed by a male mouse, and the stage of estrous for female mice. NOTE: Females that have already given birth 1-2 times will deliver larger litter sizes in future pregnancies.
1.3 Before starting timed pregnancies, house the male mouse alone in a cage one week prior to breeding. During the week that breeding initiation, introduce at least one female per male in the cage. NOTE: Depending on the approved animal protocol, adding two to three females into a breeding cage may be allowed and is preferred to increase plug generation.
1.4 Ideal mating should be arranged in the afternoon or evening before 5 pm, and females should be placed into the male’s cage or vice versa. If breeding does not occur in 4-5 days, consider switching mating partners every other day. NOTE: If unable to check a vaginal plug the following morning, separate the breeding pair the night before (i.e weekend/holidays).
1.5 Check for vaginal plugs every day in the early morning, before 10am. To check for vaginal plugs, gently lift the female mice by the base of the tail and observe the vaginal opening. Look for a white or cream-colored gelatinous mass. Often, the vaginal plug can be obvious, but if unclear, using your other hand, take tweezers and gently probe the vaginal opening. Consider only the more apparent plugs for isolation. NOTE: It is critical to check for vaginal plugs as early as possible in the morning, to avoid missing a potential pregnancy. Plugs can fall out or dissolve after 12h. NOTE: Even if a plug is observed, it does not guarantee the female mouse will be pregnant. If you observe a partial or no plug but redness near the vaginal opening of the female mice, do not consider these females for isolation as there is a less likely chance the plug will stick.
1.6 Once a vaginal plug is observed, record the day, and consider this to be E0.5. Separate the female mice from the breeding cage and plan to isolate the embryos depending on the stage of gastrulation you want to analyze.

### 2 Isolation of mouse embryos during gastrulation

2.1 Prior to starting the embryo isolation, prepare all reagents and equipment required. Clean the area thoroughly with 70% ethanol. Ensure all dissection tools (forceps and scissors) are washed and sterilized. Obtain sterile 5mL of DMEM/10% FBS and 20ml DPBS^-/-^ and place on ice. Perform all isolations using a stereomicroscope with a transmitted light stage and camera to help with gross developmental phenotyping.
2.2 Euthanize the pregnant dam by performing a cervical dislocation and start the isolation immediately. NOTE: Do not euthanize multiple pregnant dams at once as cell viability will be affected.
2.3 Place the pregnant dam on its back and sterilize the area near the vaginal opening with ethanol. Using dissection scissors and tweezers, lift the skin fold near the vaginal opening and make a small v-shaped cut, slowly reveal the uterus of the pregnant dam. Dissect out the uterine horn of the pregnant dam by holding one end of it with tweezers and cutting along it, making sure to remove the cervix. Place the uterine horn into a 10cm petri dish containing DPBS^-/-^ on ice. NOTE: Depending on the embryo isolation stage, the uterus will resemble smaller or bigger implantation sites (i.e ‘pearls’ on a string).
2.4 With dissection scissors and tweezers, cut each individual implantation site (‘pearls’) containing the decidual swellings inside and place into fresh DPBS^-/-^ in a 6cm petri dish on ice. NOTE: An average pregnant dam will have around 6-8 implanted embryos.
2.5 Take one implantation site and place it on a new 6cm petri dish on top of the stereomicroscope stage and add 500ul of DPBS^-/-^ on top of it. Adjust the focus of the microscope and the light source. NOTE: Depending on the size of the implantation site, the amount of DPBS might vary; the goal is having enough DPBS that the implantation site is submerged.
2.6 Using fine tipped dissection tweezers, remove the uterine muscle from the implantation site. Hold down the implantation site with one set of tweezers in one hand and slowly insert another pair of forceps with the other hand into the end of implantation site that was cut from the uterine horn, slowly revealing the decidua swelling. NOTE: Do not pull or tug too hard on the uterus surface, as it can lead to the rupture of the decidual swelling or even the lysis of the embryo.
2.7 After the decidual swelling is isolated, proceed to reveal the embryo. For that, hold the anti-mesometrial end of the decidual swelling with one pair of forceps and with the other pair slowly make a horizontal cut about 1/4 the size of the decidual swelling from the mesometrial end (i.e. often the more pointed end of the decidual swelling). Now with both forceps, slowly push from the anti-mesometrial end of the decidual swelling and the embryo should pop out from the freshly cut mesometrial end. NOTE: Do not tear into the decidual swelling, as this could break the embryo. If necessary, make smaller cuts along the mesometrial end of the decidual swelling.
2.8 Once the embryo is revealed, remove extraembryonic tissues attached. The parietal endodermal sac and ectoplacental cone might spontaneously come off from the embryo during the revealing process, but if not, take a pair of forceps and remove them with the associated maternal blood. Then, using two forceps, hold down the embryo with one pair and using the other set slowly peel the visceral yolk sac from the embryo. Using a P20 pipet, place the yolk sac with no more than 10μl of DPBS^-/-^ from the dish into an 8-strip PCR tube on ice, as this will be used for fast genotyping. NOTE: It is critical that the yolk sac sample is not contaminated with tissue from the pregnant dam. If this happens, it may lead to incorrect genotype assignment of the embryo.
2.9 Take bright field pictures of freshly isolated embryo to ensure staging of the littermates is similar. With a P200 pipette, slowly pipet up the embryo with 50μl of DMEM/10% FBS and place into a 1.5 ml tube on ice. The embryos will remain on ice until genotypes have been confirmed. NOTE: Embryos were kept on ice for 3-4 hours with no obvious degradation, but do not exceed this time, as a decrease in cell viability will occur. Label the embryos and genotyping tubes accordingly. Taking bright field images is encouraged to identify and annotate gross phenotypic differences among embryos.
2.10 Repeat these steps for all remaining decidual swellings. Clean all dissection tools and use new plastics for every isolation to ensure no contamination from previous isolations. NOTE: The dissection procedures should be limited to 1 hour from the moment of collection of the implantation site from the pregnant dam.

### 3 FAST Genotyping

3.1 Digest each individual visceral yolk sac in an 8-strip PCR tube. Using a P20 pipet, add 19.3μl of PCR template DNA lysis buffer and 0.7μl of 0.2μg/ml Proteinase K to each yolk sac sample. Vortex the sample and centrifuge. Place the 8-strip PCR tube in an 85°C heat block for 45 minutes and vortex every 5 minutes. NOTE: It is important to vortex samples while digesting to maximize cell lysis in 45 minutes
3.2 After 45 minutes, spin the tube strip down, and proceed with PCR for desired genetic identification. For reference, we have provided a sample protocol for a Cre-lox system. The following is an example of PCR conditions for Cre-genotyping. Design primers to amplify the 5’ and 3’ region of the targeted Cre site.
3.3 To perform the PCR reaction for cre genotyping, prepare a PCR master mix per in an 8-strip PCR tube for each yolk sac containing: 10μl Taq DNA polymerase mix, 0.5μl of 0.5μM forward Cre primer, 0.5μl of 0.5μM reverse Cre primer, 5μl PCR-certified water, and 4μl yolk sac genomic DNA. NOTE: If using a protocol that was optimized for mouse tail genotyping, add double the amount of DNA that is typically utilized for cleaner results.
3.4 Once the PCR master mix has been made, run PCR thermal cycle amplification program. For cre genotyping, the cycle is listed: (1) 95°C for 3 minutes, (2) 95°C for 30 seconds, (3) 55°C for 30 seconds, (4) 72°C for 30 seconds, Repeated step 2-4 for 34 cycles, (5) 72°C for 10 minutes, (6) 4°C hold. PCR products were run on an 1% agarose gel, and genotyping conclusions can be made. NOTE: if more than one PCR is necessary for genetic identification, run the PCR reactions simultaneously to optimize timing. NOTE: Do not consider any samples without clear genotypes. Take both the genotype and developmental stage into consideration for samples as littermates could be at different stages of gastrulation and skew results.
3.5 Store the remaining digested yolk sac samples in the -20°C freezer for long term storage.

### 4 Cell Dissociation of Embryos and Cell Viability

4.1 Once the genotypes have been confirmed, take a P200 pipet and pipet 50μl DMEM/10% FBS and pool embryos with the same genotype into a new 1.5ml tube and place on ice. NOTE: Do not proceed with the experiment if there are not at least 5 embryos per group (E7-E5) or 3 (E7.75-E8) as cell count and viability will decrease tremendously.
4.2 After embryos have been pooled based on genotypes, allow the embryos to settle to the bottom of the tube. Wash the pooled embryos by adding 50μl DPBS^-/-^. Wait for the embryos to settle to the bottom of the tube and remove as much of DPBS^-/-^ as possible without removing the embryos from the tube. Perform this step twice. NOTE: Holding the 1.5ml tube to a light source or towards a window will make it easier to see the embryos settling to the bottom of the tube.
4.3 Add 100μl of trypsin to the pooled embryos and incubate at 37°C in a heat block for 5 minutes. Gently flick the 1.5ml tubes with your fingers every 30 seconds to help the cells dissociate. NOTE: Do not vortex or use a pipet during trypsinization process as it damages the cells.
4.4 After 5 minutes, neutralize the trypsin with 300μl of DMEM/10% FBS. Centrifuge the pooled embryos at 100 g, 4 minutes, at room temperature. After centrifuging, a small pellet should appear for all samples. Resuspend the pellets in 40μl of DMEM/ 10% FBS and place on ice. NOTE: The size of the pellet will vary depending on the number of embryos pooled. It is possible that the pellet is not visible, but proceed with the next step.
4.5 Determine the concentration of the resuspended cells (40ul) using an automated cell counter. Mix 5μl cells with 5μl trypan blue in a new 1.5ml tube. Pipette mix thoroughly and pipet onto a slide to determine the cell number and cell viability. The optimal concentration of cells should be 700 to 1200 cell/μl, and the cell viability should 90% or higher. NOTE: If the concentration is lower than 200 and viability is lower than 50%, consider stopping the experiment. If cell concentration is too high, dilute the cell suspension. If cell clumps are observed, use a cell strainer to ensure single-cell suspension.
4.6 Proceed to single-cell partitioning using a microfluidic chip and follow the protocol from the microfluidic chip manufacturers^11^.

### 5 Nuclei Isolation Mouse Embryos (option for larger embryonic timepoints from E8 onward)

5.1 Prepare fresh lysis and wash buffers outlined in **Table 1** and place on ice. NOTE: Nuclei isolation can be performed on fresh or frozen samples. If samples are frozen, allow to thaw for 2-5 minutes on ice.
5.2 Prior to starting the experiment, confirm all genotypes and pool only embryos that have clear genotypes.
5.3 Using a P200, add 50μl nuclei lysis buffer to pooled embryos in a 1.5ml tube and try to pull a minimum of 3 embryos per mutant group. Allow samples to sit on ice with lysis buffer for 5 minutes and vortex every 30 seconds.
5.4 After 5 minutes of incubation, centrifuge at 500 g for 5 minutes at 4°C. Using a P200 pipet, remove the supernatant and resuspend the pellet in 50μl of wash buffer. Once resuspended, centrifuge at 500 g for 5 minutes at 4°C.
5.5 Remove the supernatant and resuspend in 40μl DPBS^-/-^. Count the cells using an automated cell counter. Mix 5μl nuclei with 5μl trypan blue in a new 1.5ml tube. Pipette mix thoroughly and load onto a slide to determine the cell number and cell viability. NOTE: All nuclei should be dyed with trypan blue; therefore, look at the % dead rather than the % live on the automated cell counter.
5.6 If the percentage of cell death is greater than 90% and the quality of nuclei looks good, continue to use a microfluidic chip, and follow protocol from the microfluidic chip manufacturers^11^.

### 6 Single-cell Partitioning (including cDNA amplification and library construction)

6.1 Proceed immediately single-cell RNA sequencing protocol following the microfluidic chip manufacturers protocol^11^, for the most optimal results set target cell recovery to 6,000 cells or greater.

### 7 Sequencing

7.1 After libraries are constructed, the fragment size distribution and concentrations of samples are measured using an automated electrophoresis analyzer. NOTE: Average fragment sizes distributions should be between 300-1,000bp.
7.2 Samples are ready to be loaded into the sequencer. In a standard sequencing protocol libraries from different samples are pooled together. The final loading concentration of pooled libraries should be 750pM in a total volume of 24μl. Load pooled libraries into reagent cartridge, following sequencing manufacturer’s protocol^12^.
7.3 Libraries loaded into the pre-assembled flow cell and cartridge are sequenced according to the desired sequencing depth with pair-end, dual indexing. The sequencing reads adhered to the protocol outline by microfluidic chip manufacturers is as follow^11^: Read 1: 28 cycles, i7 Index: 10 cycles, i5 Index: 10 cycles, and Read 2: 90 cycles. Following the completion of sequencing, the data undergoes bioinformatics analysis.
7.4 Use bioinformatics methods to perform quality controls. This includes the number of sequenced cells, reads per cell, and number of genes mapped per cell. NOTE: For optimal results, the number of sequenced cells should be at least the 80% of targeted cells. It is recommended that the number of reads per cell is at least 30K and the number of genes detected higher than 3000 (for mouse samples).

## REPRESENTATIVE RESULTS

The methodology designed in this paper is specifically designed to enhance the preparation of embryo samples for single-cell omics from E6.5 to E8. This robust pipeline consists of five major steps: synchronized timed pregnancies, embryo isolations, FAST genotyping, cell dissociation, and cell viability **(Figure 1A)**. While the presented data focuses on timepoints from E7 to E7.5, it can be applied up to E8 embryos **(Figure 1B)**. Synchronized timed pregnancies were performed by placing two female mice into a cage with a male mouse. Vaginal plugs were checked every morning before 10AM, and if a plug was observed, it was considered E0.5 by noon of that day **(Figure 2A)**. Optimal conditions for plugs required a female in estrus or proestrus stage paired with a male having a history of successfully placing several plugs in prior breeding **(Figure 2B)**. Only obvious plugs were considered for embryo isolations **(Figure 2C)**.

**Table 1:**
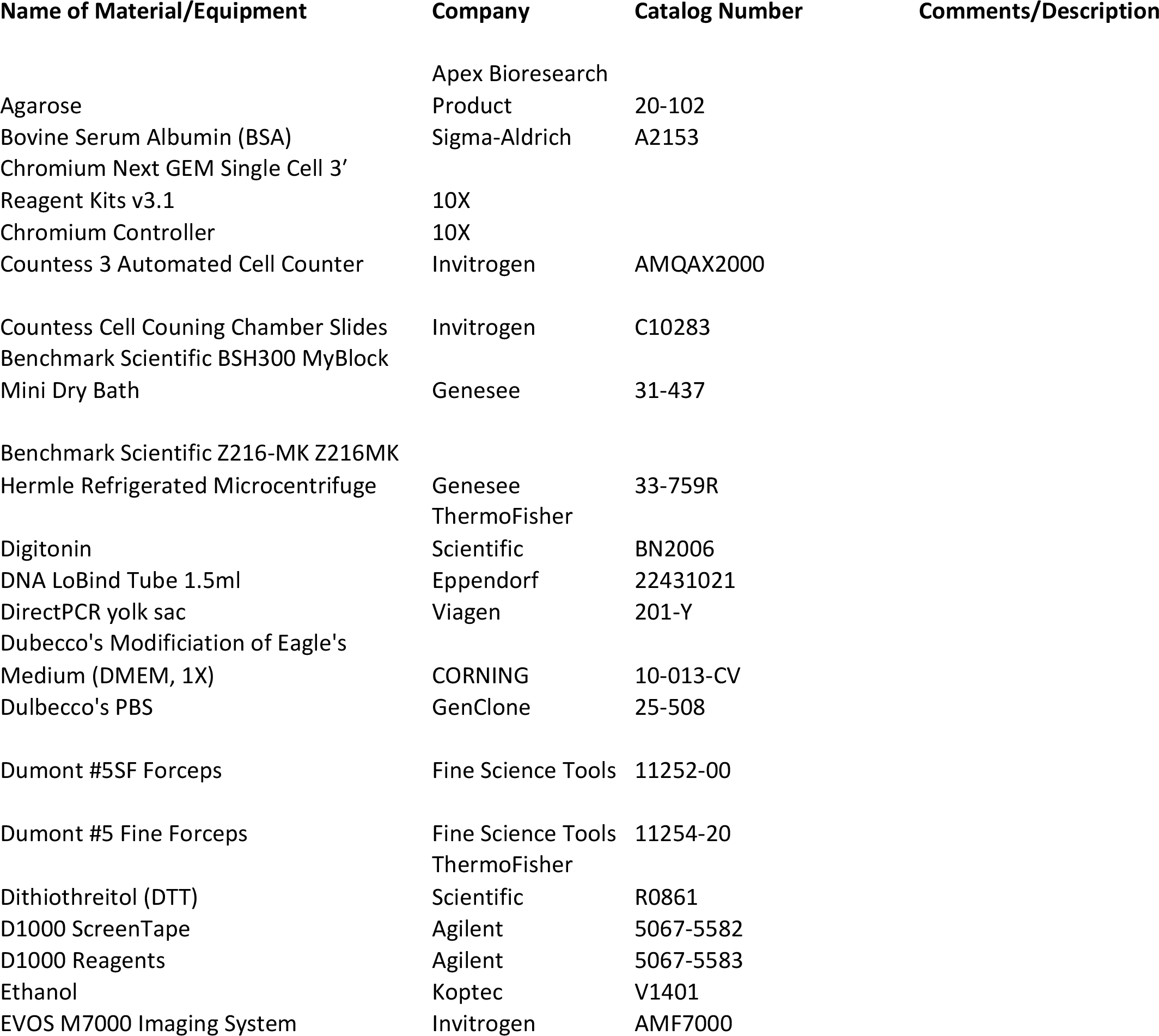
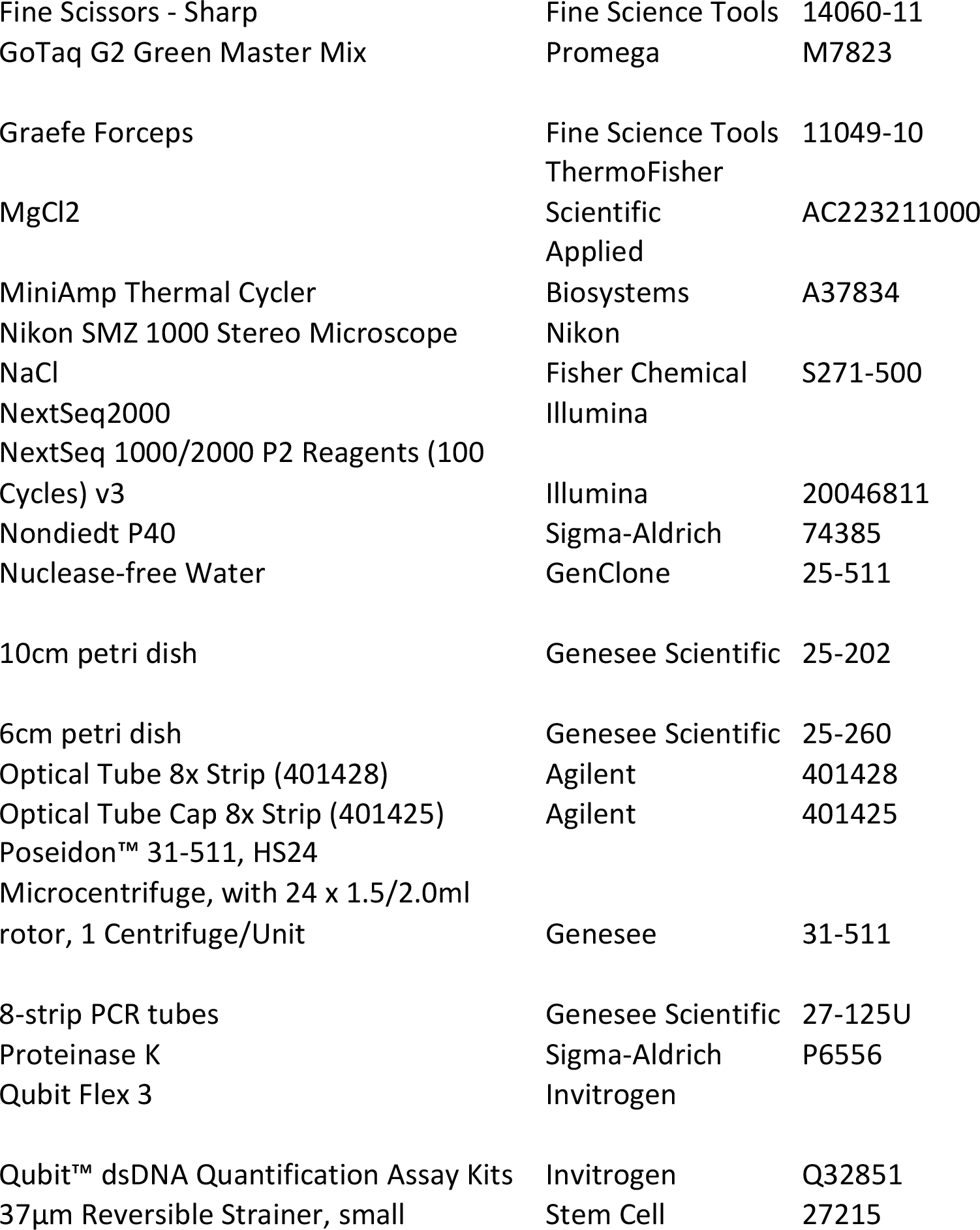
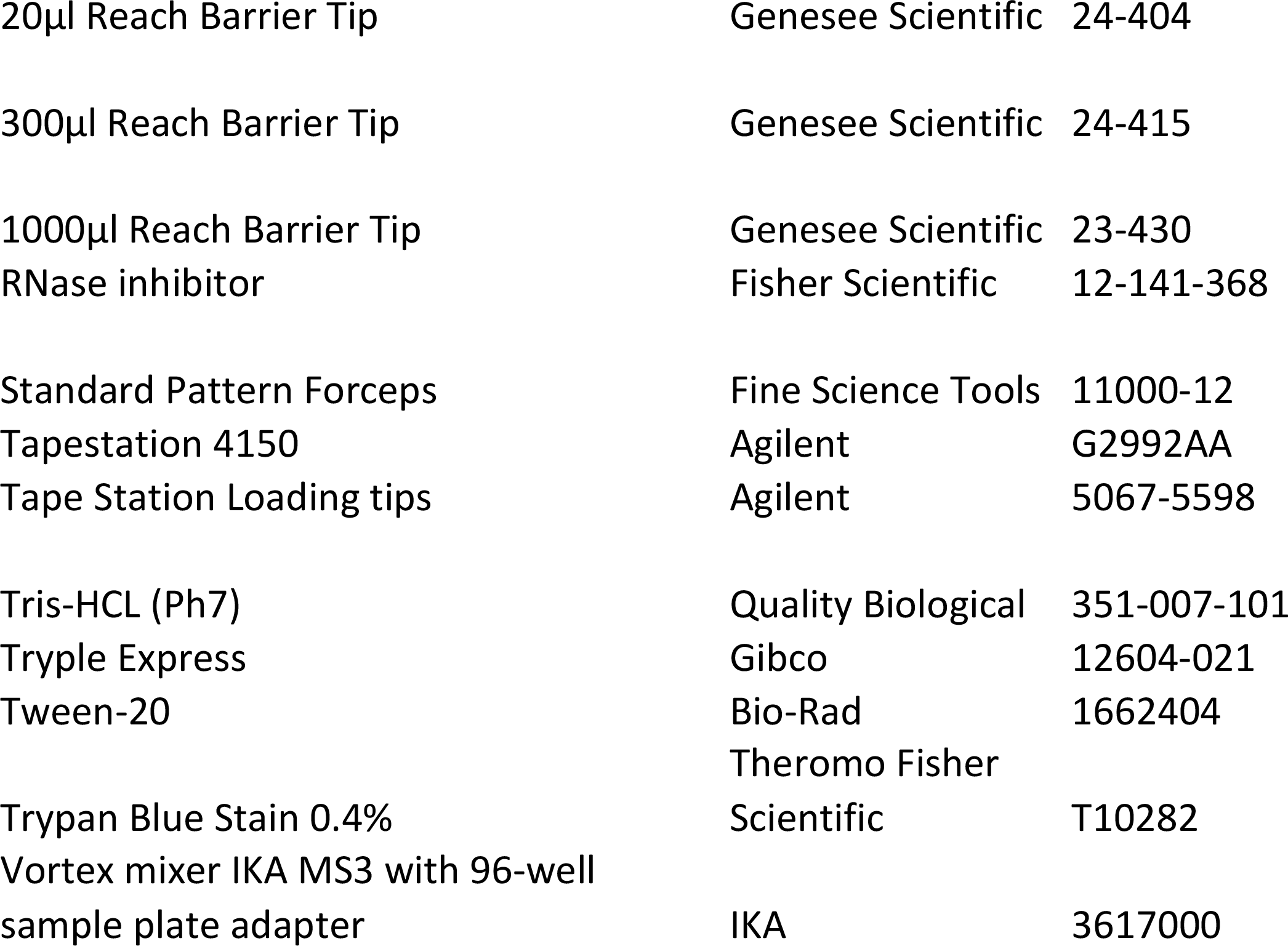
Nuclei Isolation Lysis Buffer.

**Figure 1.**
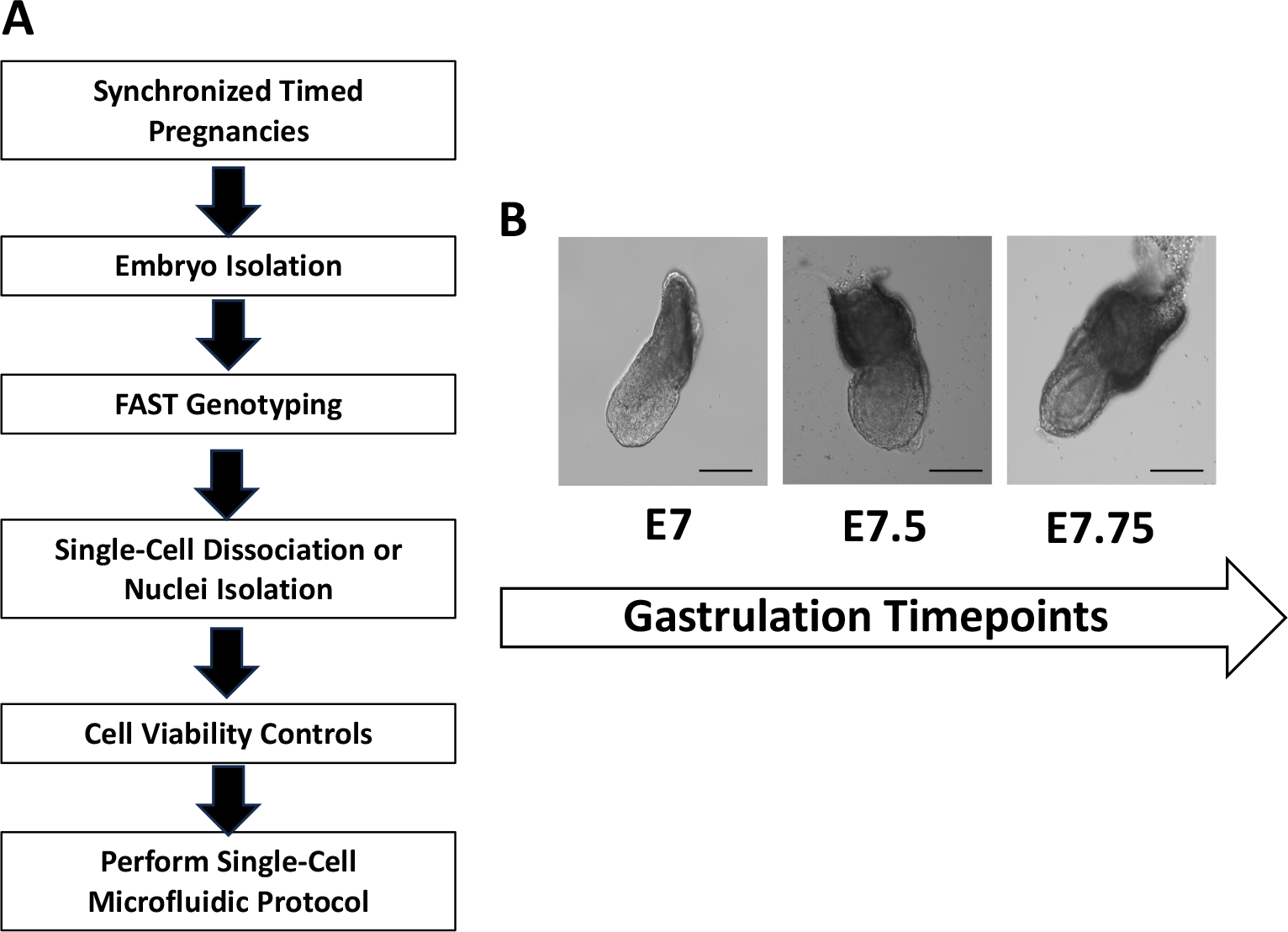
Optimization of early gastrulating whole mouse embryos for single-cell RNA sequencing. (**A**) Workflow schematic for obtaining high-quality cells and/or nuclei from gastrulating embryos **(B)** Representative bright field images of mouse embryos during gastrulation from E7 to E7.75. Scale bar, 125 μm

**Figure 2.**
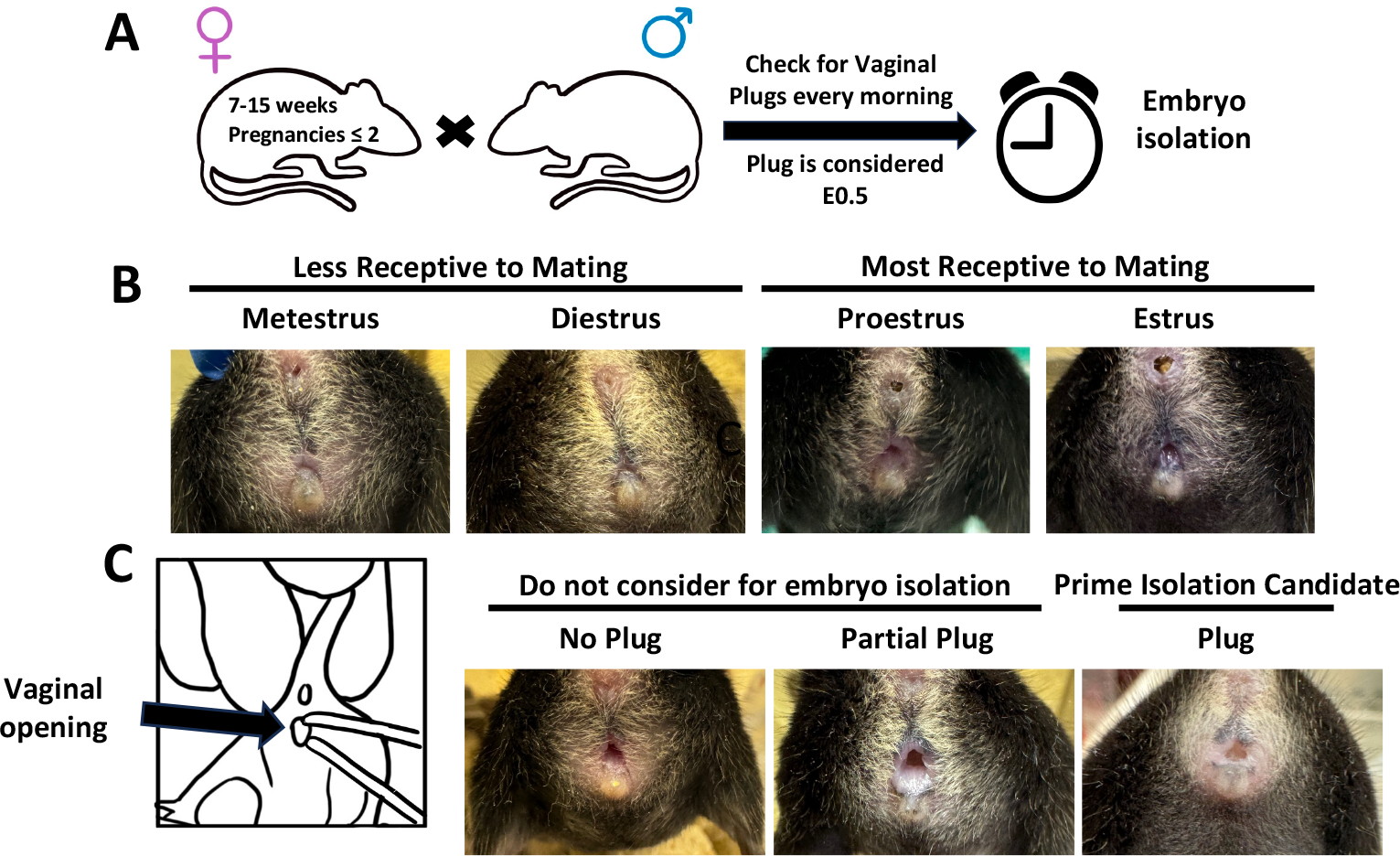
Strategy for efficient synchronized timed pregnancies for embryo isolations during gastrulation. **(A)** Schematic diagram indicating the pipeline for timed pregnancies **(B)** Representative images of the phases of estrus cycle for female mice. The phases most receptive to breeding are indicated **(C)** Schematic diagram illustrating how to check for vaginal plugs and examples of vaginal openings without a plug, a partial plug or a good plug to consider for embryo isolations.

**Figure 3** exemplifies a successful embryo isolation at E7.5. We isolated the uterine horn, the decidual swelling was individually cut, the embryo was revealed, and the yolk sack was isolated for genotyping. FAST genotyping was performed within 3 hours of embryo isolation **(Figure 4A)**. The visceral yolk sack, the parietal endodermal sac and ectoplacental cone with associated maternal blood is shown in **Figure 4B & C**. The visceral yolk sac was utilized for genotyping **(Figure 4C)**. After digesting the yolk sac, the PCR mix was prepared, and the PCR reaction was run. The resulting PCR product was then separated on an agarose gel. We present an example of FAST genotyping using a Cre-recombinase system. The expected fragment sizes for our system in flox genotyping were 597 bp for flox and 498 bp for the wild type. The expected fragment size for Cre genotyping is 650 bp **(Figure 4D)**. Representative gels for Cre and flox are shown in **Figure 4E**.

**Figure 3.**
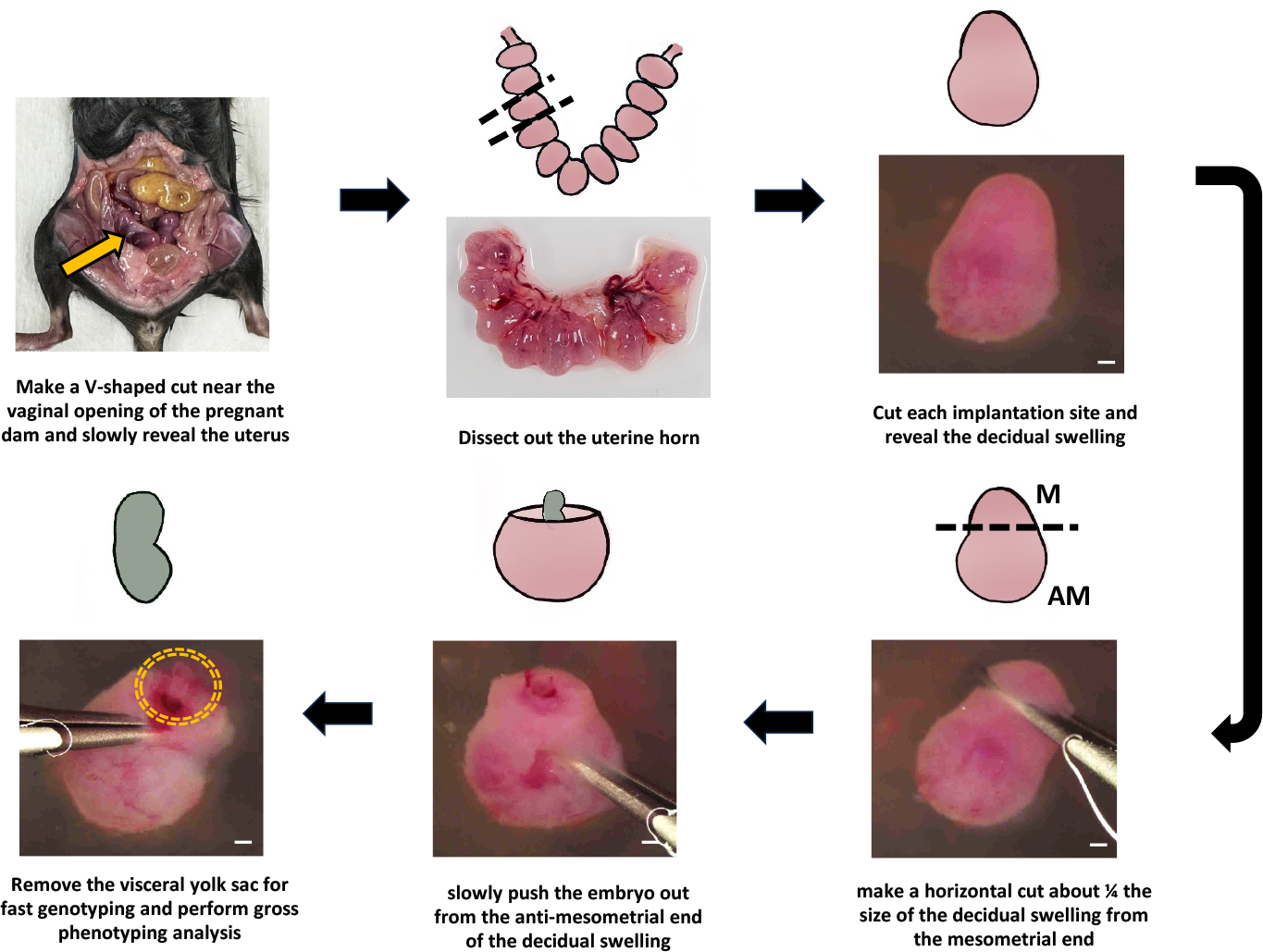
Dissection and genotyping of E7.5 embryos for single cell RNA sequencing. Schematic diagrams and images of the process of isolating E7.5 embryos and the dissection of the visceral yolk sac. The yellow arrow indicates the uterus of the pregnant dam, and the yellow dashed circle outlines the location of the embryo. Dashed lines represent the areas that were cut during dissection. M means mesometrial end, while AM means anti-mesometrial end. Scale bars from stereoscope images are 400μm.

**Figure 4.**
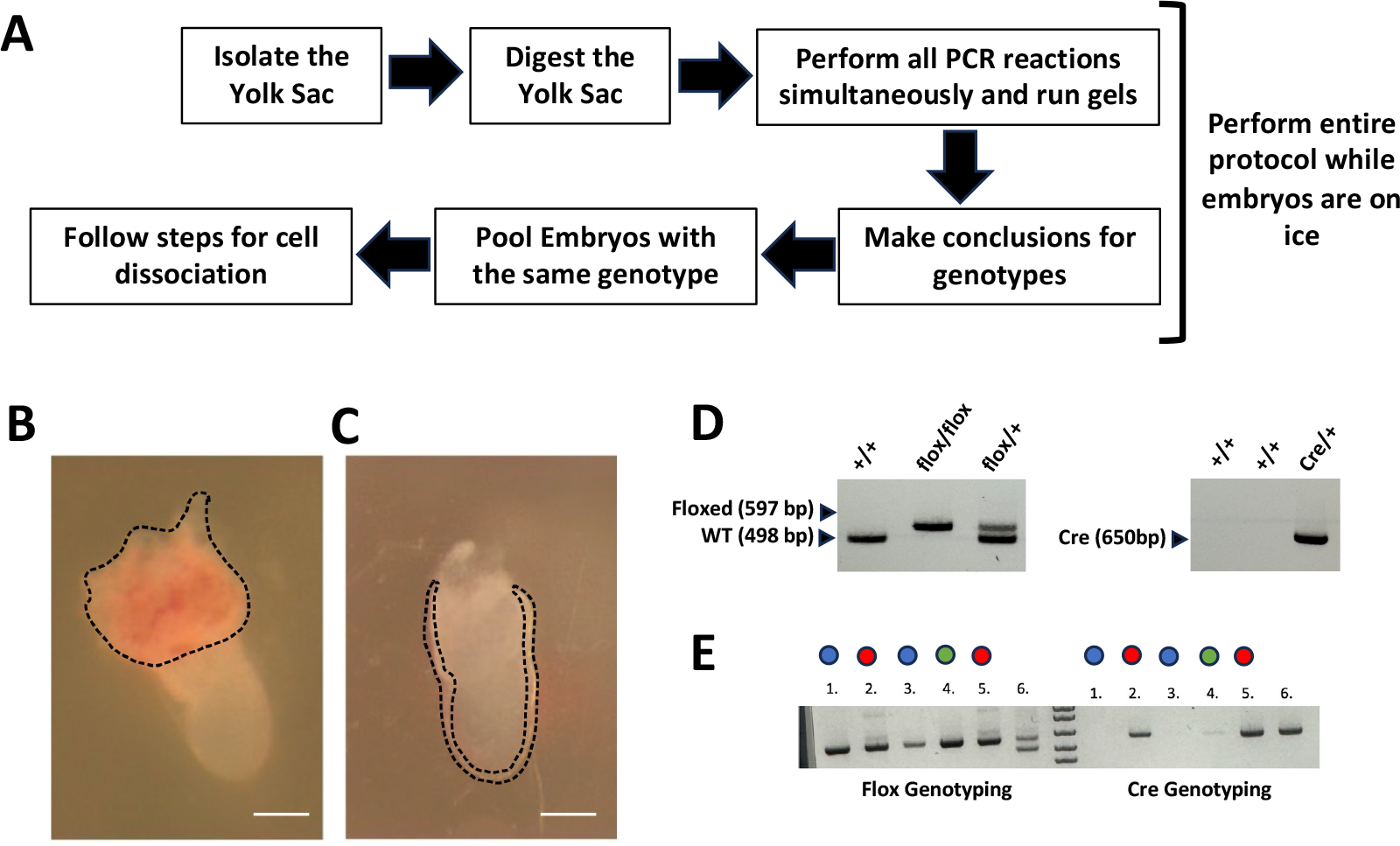
FAST genotyping of yolk sacs from gastrulating mouse embryos. **(A)** Workflow schematic for same-day genotyping of yolk sacs **(B)** Representative image of the parietal endodermal sac and ectoplacental cone with associated maternal blood **(C)** Visceral yolk sac in E7.5 embryos highlighted by dashed line, scale bar, 200μm **(D)** Representative gels of wild type, heterozygous, and homozygous flox and Cre genotyping with DNA fragment sizes **(E)** FAST genotyping results for Cre and flox genotyping from yolk sacs, 2 controls indicated in blue and 2 mutant embryos indicated in red were pooled and processed for single cell RNA sequencing. 1 embryo indicated in green shows an unclear genotype and was not used for further processing.

Cell count and good viability are required for a successful single cell experiment. Suspensions with low cell viability, high percentage of dead cells, clumping, or high debris are unsuitable for further processing. Optimal conditions are for a 700-1,200 number of live cells per μl and >90% viability. **Figure 5A** presents a panel illustrating both good and sub-optimal cell viability. The same criteria can also be applied for nuclei isolation, as shown in **Figure 5B**. However, it is crucial to note that the evaluation of trypan blue differs from cells and nuclei: viable cells do not incorporate trypan blue while viable nuclei do. If cells/nuclei suspension are optimal (>90% viability), proceed with single-cell partitioning using a microfluidic chip following manufacturers procedures^11^. Troubleshooting options are provided for suspensions where cell/nuclei viability is between 60-89%, as depicted in **Figure 5C**. If viability, regardless of the total number of cells, falls below 60% consider stopping the experiment.

**Figure 5.**
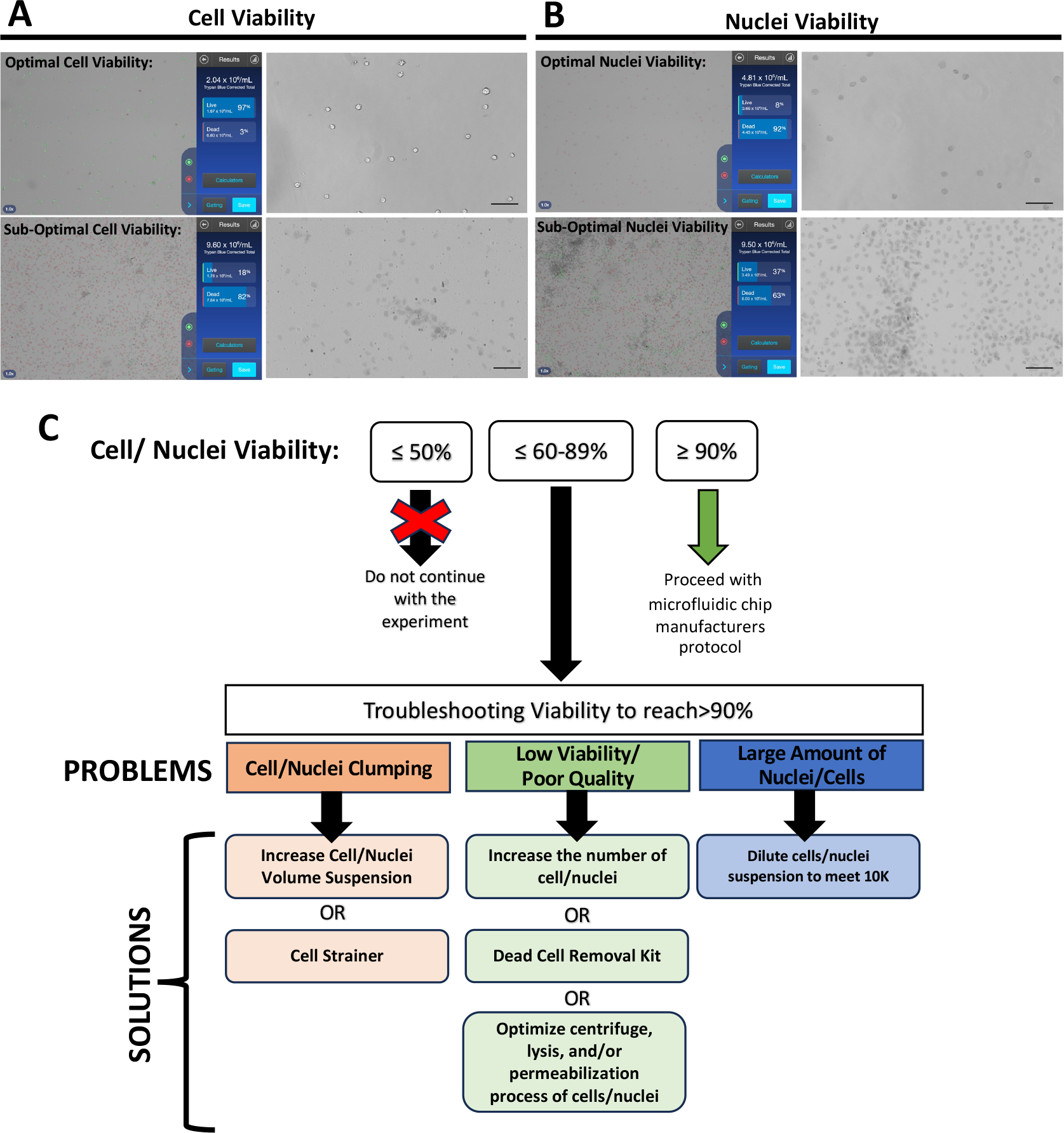
Assessment of cell quality and nuclei viability. **(A)** Representative images of optimal and sub-optimal cell viability conditions from E7.5 embryos **(B)** Representative images of optimal and sub-optimal nuclei viability conditions from E8 embryos **(C)** Troubleshooting scheme indicating potential solutions to help increase the viability of cells before starting single-cell partitioning.

Following the procedures outlined by the single cell manufacturers procedures^11^, library constructions were prepared for single-cell RNA sequencing using cells obtained from E7 embryos, with cell viability ranging from sub-optimal to optimal conditions, aiming for an estimated target recovery of 2,000 cells **(Figure 6). Figure 6A** depicts a representative fragment size distribution of scRNAseq libraries for both sub-optimal and optimal conditions, indicating that cell viability does not significantly affect the entire single cell partitioning process. The fragment sizes distribution ranged between 400-500bp. This indicates that sub-optimal conditions do not affect the process of library preparation. **Figure 6B** shows the outcomes of both successful and sub-optimal scRNAseq experiments. Following sequencing, we conducted quality control checks on samples and observed that, in cases where cell viability was sub-optimal, only 10% of cells were successfully sequenced. In contrast, optimal samples exhibited a higher percentage, with 91% of the total cells being sequenced. This is further proven by barcode plots, indicating that in the sub-optimal conditions have larger background noise compared to optimal conditions. Clustering analysis was performed for both conditions and shows that 9 clusters were revealed in the optimal conditions and only 4 were revealed in the sub-optimal, highlight the importance of high-quality cells required for proper representation of data during these stages of development.

**Figure 6.**
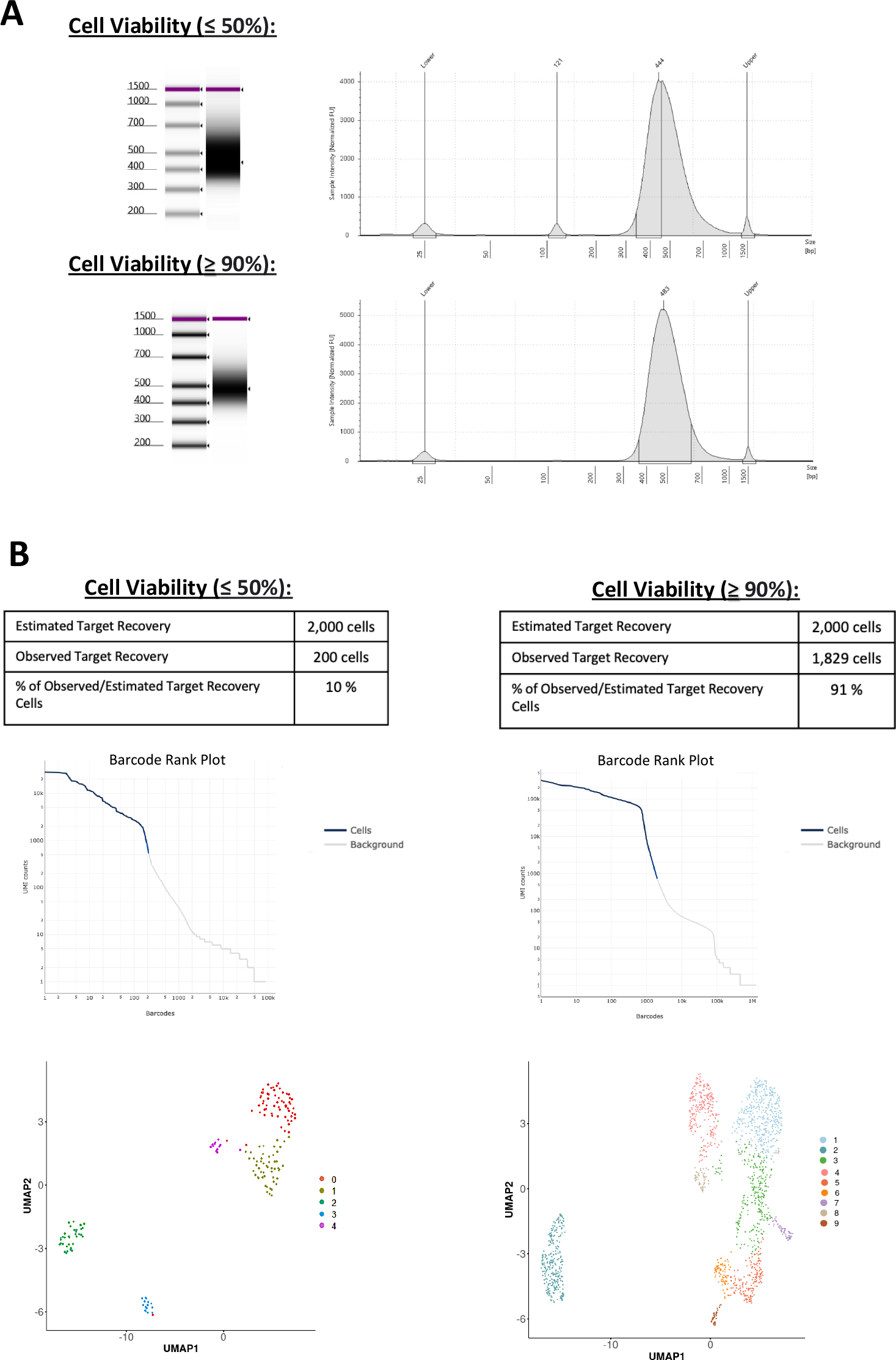
Single cell RNA sequencing of E7 mouse embryos. **(A)** Representative trace of fragment size distribution for single cell RNA sequencing libraries from E7 mouse embryos for both sub-optimal and optimal conditions of cell viability, with the main peak near 400-500 bp **(B)** Analysis of single cell RNA sequencing outcomes for E7 mouse embryos under both sub-optimal and optimal conditions showing observed target recovery, barcode ranking, and clustering of cell types through U-MAP distribution.

## DISCUSSION

We present here a robust pipeline for obtaining high-quality single-cell and nuclei suspensions from gastrulating mouse embryos, specifically designed to facilitate studies on mechanisms of cell-fate specification in early development. This method addresses a crucial gap in the field of gastrulation, optimizing the analysis of embryos requiring genotypes, such as sex or somatic genes. Utilizing genetic mutation mouse models and employing high-resolution single cell sequencing on whole mouse embryos, this pipeline can further enhance the understanding of the gene expression profiles of the early mouse gastrula. We demonstrate the feasibility of using a genetic mutation mouse model by a Cre recombinase system, obtaining high quality cells and nuclei at early timepoints of development for single cell -omics. This method evolved after multiple attempts, during which our samples did not meet the quality standards required for library preparation and sequencing. The explained methodology generates sufficient cells/nuclei from <E8 embryos through optimization of three critical steps: 1) synchronized timed pregnancies to increase the number of embryos 2) FAST genotyping to avoid freezing/thawing, and 3) assessing cell/nuclei viability to avoid sequencing of dying cells.

At the E7 embryo, each embryo typically consists of around 300 cells and obtaining embryos with the desired genotype is challenging, with only 1 or 2 embryos per pregnant dam meeting the criteria. Attempts to increase embryo pool size by snap-freezing embryos from pregnant dams on different days proved unsuccessful, as cell viability was severely compromised after thawing, even with snap-freezing or the addition of cryopreserving agents. To address this challenge, we optimized the breeding strategy and synchronized pregnant dams. To increase the chances of multiple isolations, we recommend using female mice that have given birth 1-2 times before breeding for isolation, as they are more likely to have larger litter sizes. Monitoring the female’s estrous cycle is crucial; mating is more likely to occur during the proestrus and estrus stages. If breeding difficulties arise, switching the breeding partners after four days could also help if no plug is produced. It’s important to note that this method has a limitation: it relies on observed plugs and even if a plug is observed, it does not guarantee pregnancy, just indicates sexual activity. Therefore, increasing the number of plugs in a day will increase the probability of more than one pregnant dam in a day and number of positive genotypes.

Having good cell viability is essential for the success of single-cell sequencing. In the microfluidic chip design provided by the manufacturers, single cells are partitioned into Gel Beads-in-emulsion (GEMs) within a chip containing known barcoded gel beads^11^. However, a notable limitation of this process is that both high-quality and poor-quality single cells can be partitioned. Even with an adequate number of live cells (i.e. 1,000) if the viability of the suspension is low (i.e. 1000 cells alive and 1000 cells dead, resulting in 50% viability), the sequencing experiment will likely fail. For optimal results, we recommend aiming for a viability around 90%. If the cell viability falls between around 60-89 %, specific measures can be taken to enhance the experiment’s viability. However, if the cell viability is less than 60%, we strongly advise against continuing with the experiment. The reason for this is that the dying cells will be “captured” in the partitioning gel and subsequent library preparation and quality controls will pass without noticeable issues. However, the actual sequencing experiment may completely fail as most of the sequencing reads will map mitochondrial, ribosomal or apoptosis genes, indicating signs of poor cell viability from the outset. Our data illustrates a pipeline for single cell sequencing of gastrulating mouse embryos with high quality cells. This methodology can be implied for the understanding of many different genetic mutations mouse models during development.

## ACKNOWLEDGMENTS

We acknowledge the Genomics Core at the Fox Chase Cancer Center and Dr. Johnathan Whetstine laboratory for technical support for the sequencing experiments. We acknowledge laboratory members of Dr. Estaras, and Alex Morris, a rotation graduate student who contributed to the initial analysis of the single-cell studies. This work is funded by the NIH grants R01HD106969 and R56HL163146 to Conchi Estaras. Additionally, Elizabeth Abraham was supported by T32 training grant 5T32HL091804-12.

